# Mutational fitness landscapes of diverse human influenza H1N1 neuraminidases

**DOI:** 10.64898/2026.10.07.757144

**Authors:** Qi Wen Teo, Zongjun Mou, Wenkan Liu, Huibin Lv, Kevin J. Mao, Jessica J. Huang, Yang Wei Huan, Ruipeng Lei, Akshita B. Gopal, Arjun Mehta, Emily X. Ma, Xinghong Dai, Nicholas C. Wu

## Abstract

While neuraminidase (NA) is emerging as a promising influenza vaccine target, it evolves rapidly and undergoes antigenic drift. However, our understanding of its evolutionary constraints remains limited. Here, we systematically quantified the replication fitness effects of >6,000 mutations for each of three antigenically diverse H1N1 NAs using deep mutational scanning. Moderate correlations among H1N1 NA fitness landscapes revealed mutations with strain-dependent replication fitness effects. Additionally, cross-subtype comparison between N1 and N2 NAs uncovered similar epistatic effects that were enriched in the C-terminal interprotomer interface. We also showed that NA mutational fitness strongly correlated between *in vivo* and *in vitro*. Characterization of 40 individually constructed NA mutants further demonstrated that NA mutational fitness depended more on its surface protein expression than on its total protein expression or endoplasmic reticulum stress induction. Overall, the large dataset generated in this work provides critical insights into the evolutionary constraints of NA.

## INTRODUCTION

Influenza A virus (IAV) remains a major global public health threat due to its continuous evolution and capacity to cause seasonal epidemics and pandemics^1^. IAV has two surface glycoproteins, hemagglutinin (HA) and neuraminidase (NA). To date, 19 HA subtypes (H1-H19)^2^ and 11 NA subtypes (N1-N11) have been identified^3^. Among these, H1N1 and H3N2 are the two subtypes currently circulating in the human population and are responsible for seasonal influenza epidemics^4^. HA initiates virus entry by engaging sialylated glycan receptors, whereas NA facilitates progeny virion release by cleaving sialic acids^5^. While HA is traditionally the focus of influenza vaccine development, NA has emerged in recent years as an effective vaccine target^6^. NA is also a target for three FDA-approved inhibitors, zanamivir, oseltamivir, and peramivir. As a result, the evolution of NA has important implications for antigenic drift and drug resistance.

NA is a homotetrameric type II transmembrane glycoprotein composed of an N-terminal cytoplasmic tail, a transmembrane domain, a stalk domain, and a C-terminal globular head domain that contains a highly conserved enzymatic active site^7^. Tetramerization of NA appears to be critical for its folding stability, since NA enzymatic activity and immunogenicity are much higher in the tetrameric form than in the monomeric form^7,8^. The NA active site comprises eight catalytic residues that interact directly with sialic acids and 11 framework residues that maintain the structural conformation of the active site^7^. The globular head domain also contains major antigenic sites surrounding the active site^9^. Consistently, antigenic drift of human NA is mainly driven by mutations adjacent to the active site^10–12^.

The emergence and continued circulation of NA inhibitor resistance mutations remain a significant challenge for influenza control^13^. For instance, H274Y, which confers strong resistance to multiple NA inhibitors, is readily detected in circulating H1N1 IAVs^13^. Similarly, N294S, which is identified in both N1 and N2 subtypes, is associated with reduced sensitivity to oseltamivir^14^. Along with antigenic drift of NA, these observations highlight the evolutionary plasticity of NA and underscore the need to better understand its underlying constraints. While we have previously mapped the mutational fitness landscape of H3N2 NA and also demonstrated the impact of epistasis on its evolution^15,16^, little is known about the evolutionary constraints of H1N1 NA. Notably, N1 and N2 subtypes belong to group 1 and 2 NAs, respectively, and hence are phylogenetically distant.

Here, we performed deep mutational scanning to measure the fitness effects of nearly all possible single amino acid mutations in the NA head domain of three different influenza H1N1 strains. Our results showed that the mutational fitness landscapes among different N1 and N2 NAs were moderately correlated, which led to the identification of mutations with strain-dependent replication fitness effects. We have also demonstrated that the replication fitness effects of NA mutations were largely determined by their protein surface expression. Moreover, NA mutational fitness landscape was strongly correlated between *in vitro* and *in vivo*.

## RESULTS

### Mutational fitness landscape of N1 NA is similar *in vitro* and *in vivo*

To systematically evaluate the fitness effects of H1N1 NA mutations, we performed deep mutational scanning of H1N1 A/Puerto Rico/8/1934 (N1/PR8) NA both *in vitro* and *in vivo*. Briefly, saturation mutagenesis was applied to generate all possible single amino acid mutations spanning residues 82-470 (N2 numbering used throughout) in N1/PR8 NA, covering 99% of the head domain. The virus mutant library was rescued and passaged in MDCK-SIAT1 cells (hereafter referred to as P1 library), followed by an additional passage in MDCK-SIAT1 cells (hereafter referred to as P2 library) or in mice (hereafter referred to as mouse library) (**Figure S1**). Subsequently, next-generation sequencing was used to quantify the frequencies of individual mutations in the plasmid and post-passaged mutant libraries, which in turn were used to calculate their fitness scores. The average fitness scores of silent mutations and nonsense mutations were normalized to 1 and 0, respectively. We measured the fitness scores of 7,072 (97%) of 7,296 all possible amino acid mutations in the N1/PR8 NA head domain, both *in vitro* and *in vivo* (**Figure 1 and Table S1**). A Pearson correlation of 0.76, 0.77 and 0.85 for the P1, P2 libraries and mouse library were obtained between biological replicates (**Figure S2A-C**), demonstrating a high reproducibility of our deep mutational scanning result. Furthermore, mutational fitness scores strongly correlated between P1 and P2 libraries (R = 0.96) and between P2 and mouse libraries (R = 0.94) (**Figure 2A and Figure S2D-E**), indicating that the selection pressures in cell culture and in mice were similar. Of note, fitness scores from P1 library were used for all the downstream analysis.

**Figure 1.**
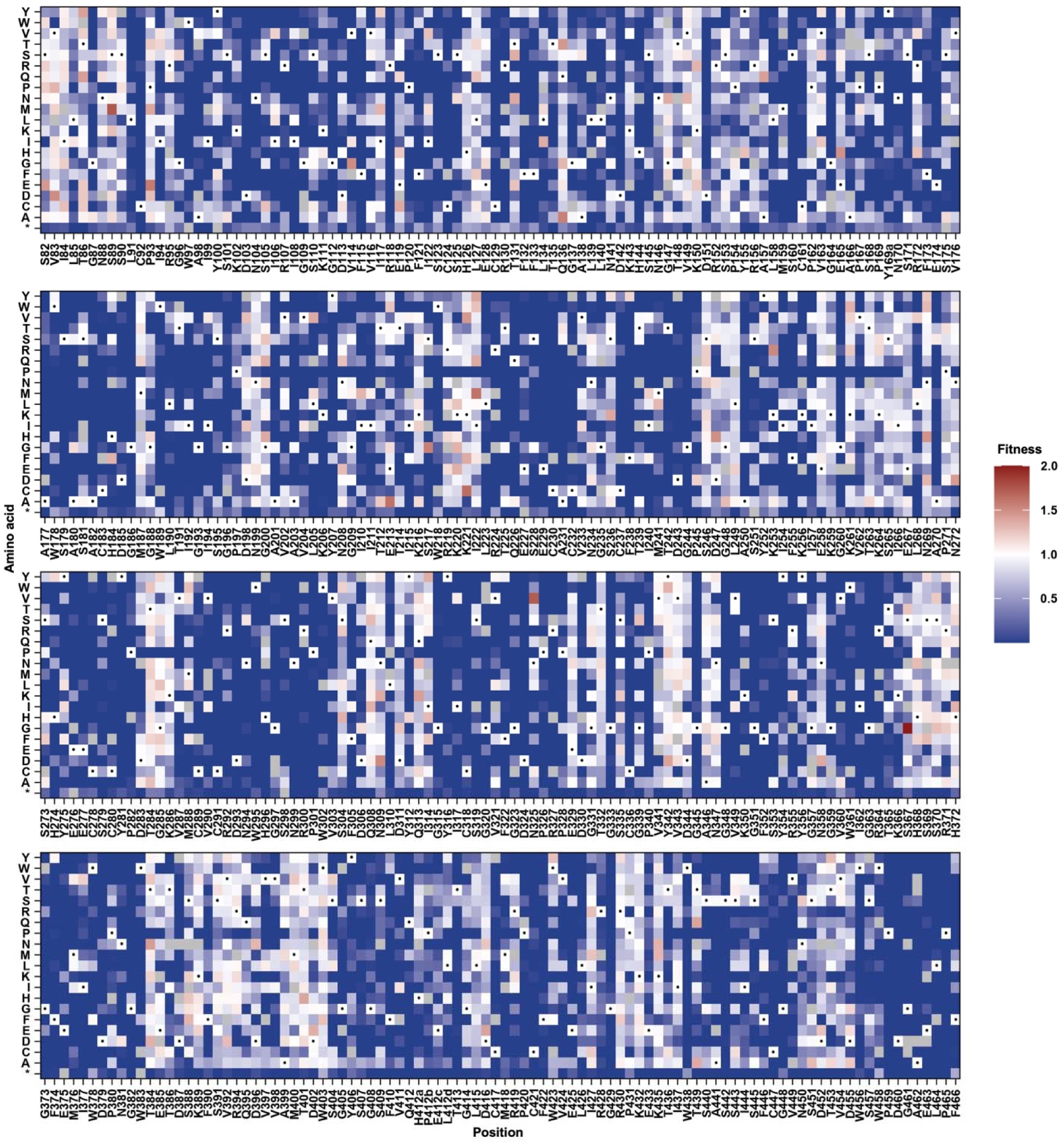
Deep mutational scanning of N1/PR8 NA. The replication fitness effects of individual mutations in the N1/PR8 NA P1 library were measured by deep mutational scanning and the fitness scores are shown as a heatmap. Wild-type (WT) amino acids are indicated by a black circle. Mutations in gray were excluded in our data analysis due to low input count. Stop codon is annotated as *.

**Figure 2.**
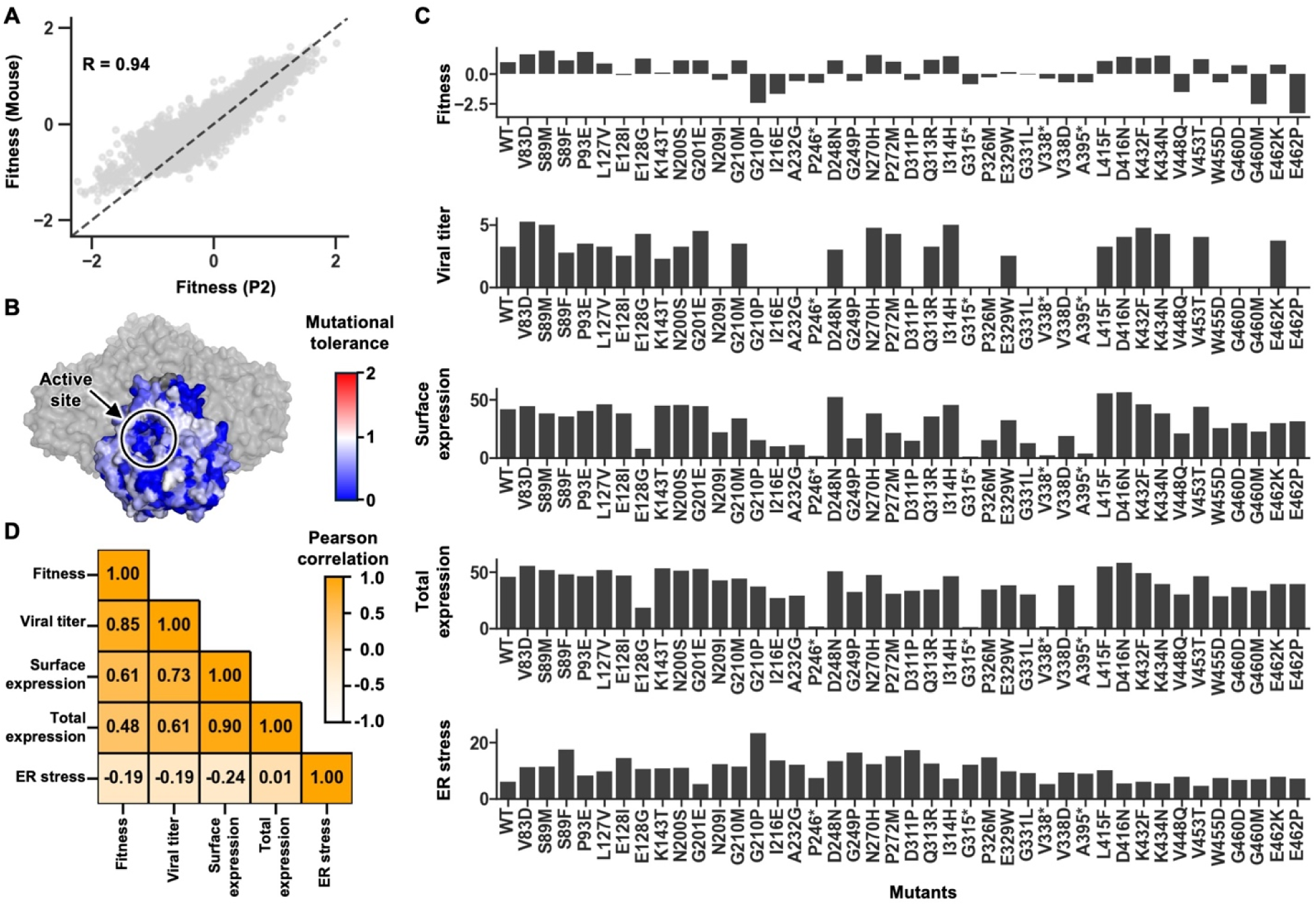
Functional consequences of N1/PR8 NA mutations on expression, trafficking, and ER stress. **(A)** Scatterplot showing the correlation of fitness scores for individual mutations between N1/PR8 P2 and mice library. Pearson correlation coefficient (R) is indicated. **(B)** Mutational tolerance across NA residues of N1/PR8 P1 library mapped onto the NA structure from H1N1 A/Brevig Misson/1/1918 (PDB: 8E6K)^35^. One protomer is displayed in surface representation with residues colored according to mutational tolerance, while the remaining three protomers are shown in gray. Active site is circled. **(C)** Bar charts showing the fitness score, viral titer, NA surface expression, NA total expression and ER stress for 40 mutants. **(D)** Pearson correlation coefficients among fitness score, viral titer, NA surface expression, NA total expression and ER stress are shown as a heatmap.

We further quantified mutational tolerance for each residue based on the average fitness scores of all mutations at a given residue. Mapping mutational tolerance onto the NA structure revealed strong evolutionary constraints within or proximal to the catalytic pocket (**Figure 2B**). Catalytic residues in NA active site, R118 and Y406, and the adjacent framework residues, W178, S179, E277, and E425, were highly intolerant to mutation (**Figure S2F**), consistent with the absence of known NA inhibitor resistance mutations at these positions. In contrast, among all mutations at H274 and N294, only the clinically relevant resistance mutations H274Y and N294S showed no fitness cost (**Figure 1 and Table S1**). Residues with high mutational tolerance were also identified within the 150- and 430-loops, including G147, V149, K150, Q430, and K432, which are immediately proximal to the active site (**Figure S2F**). Together, these observations illustrated that our deep mutational scanning data is informative for understanding the evolutionary constraints of H1N1 NA.

### Surface NA expression is a key determinant of viral replication fitness

We have previously demonstrated that IAV replication fitness is influenced by NA protein expression and surface trafficking^17^. Given that NA expression can also induce endoplasmic reticulum (ER) stress^18^, we aimed to determine how NA protein expression, NA trafficking, and NA-induced ER stress collectively shaped the replication fitness. To this end, we individually introduced 40 mutations into N1/PR8 NA, including 36 missense mutations spanning a broad range of fitness scores from our deep mutational scanning experiment (–3.35 to 1.95) and four nonsense mutations as negative controls (**Figure 2C**). The impacts of each of these 40 mutations on viral titer, total NA expression, surface NA expression, and ER stress induction were measured. A strong correlation between viral titer and fitness scores from the deep mutational scanning result was observed (R = 0.85) (**Figure 2D and S2G**). Further analysis revealed that viral titer correlated stronger with surface NA expression (R = 0.73) than with total NA expression (R =0.61) (**Figure 2D and S2G**). In contrast, ER stress induction had a weak negative correlation with viral titer (R = −0.19) and with surface NA expression (R = −0.24), while showing almost no correlation with total NA expression (R = 0.01) (**Figure 2D and S2G**). These data illustrated that the replication fitness effects of NA mutations were primarily determined by their impacts on their surface protein expression levels.

### Comparing NA mutational fitness landscapes between seasonal human H1N1 strains

Given that N1/PR8 is a mouse-adapted strain, we further performed deep mutational scanning experiments on the head domains of two seasonal human H1N1 NAs, namely A/Marton/1943 (N1/Mar43) and A/Georgia/20/2006 (N1/Geo06). These two seasonal human H1N1 strains were isolated 63 years apart and their NAs were differed by 52 amino acid mutations. Additionally, N1/Geo06 contains a Thr insertion at position 435 (T435a) that emerged in human H1N1 viruses during late 1940s and disappeared following the 2009 H1N1 pandemic. Our deep mutational scanning experiments quantified the replication fitness effects of 92% and 97% of all possible amino acid mutations in the NA head domains of N1/Mar43 and N1/Geo06, respectively (**Figure S3-S4 and Table S1**). A Pearson correlation of 0.55 and 0.47 was obtained between two biological replicates for Mar43 and Geo06 libraries (**Figure S5A-B**), demonstrating the reproducibility of our deep mutational scanning result. To compare the mutational tolerance of NA between N1/Mar43 and N1/Geo06, their difference was projected onto a cryogenic electron microscopy (cryo-EM) structure of N1/Mar43 NA that was determined in this study at a resolution of 1.82 Å (**Table S2**). While most NA residues had similar mutational tolerance between N1/Mar43 and N1/Geo06, some residues exhibited substantial difference, including the C-terminal residues A462, E463, L464, and P465 (**Figure 3A**). Similar observations were made when comparing the NA mutational tolerance of N1/Mar43 and N1/Geo06 to that of N1/PR8 (**Figure S5C-D**). Consistently, the NA mutational fitness effects correlated only moderately between N1/Mar43 and N1/Geo06 (R = 0.41), between N1/PR8 and N1/Mar43 (R = 0.52), and between N1/PR8 and N1/Geo06 (R = 0.59) (**Figure 3B and S5E-F**). These results suggested that epistasis played a role in shaping the mutational fitness landscape of H1N1 NA.

**Figure 3.**
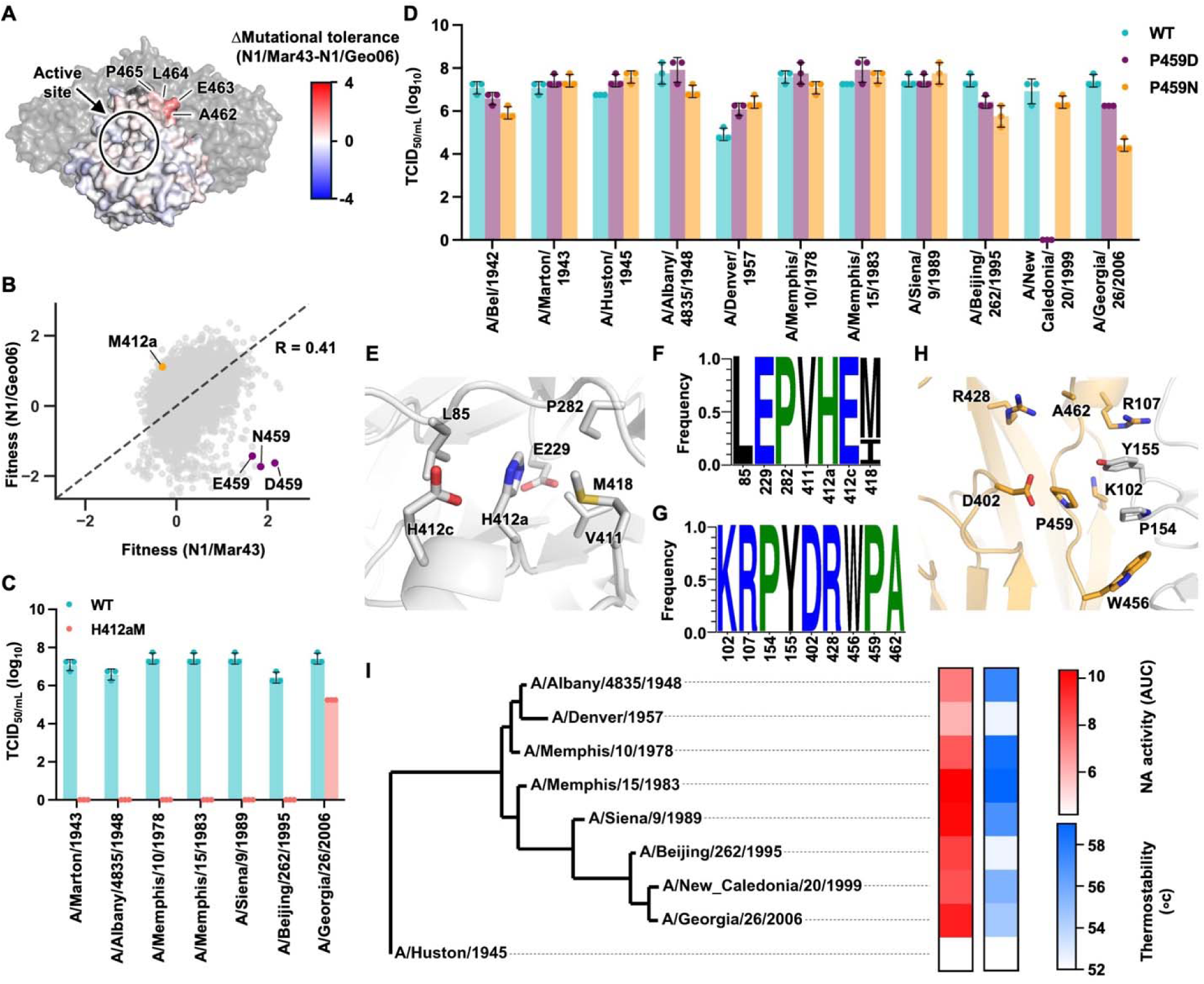
Cross-strain comparison of NA mutational effects in human H1N1 viruses. **(A)** Difference of mutational tolerance between N1/Mar43 and N1/Geo06 mapped onto the cryo-EM structure of N1/Mar43 NA. One protomer is displayed in surface representation with residues colored according to mutational tolerance, while the remaining three protomers are shown in gray. Active site is circled. **(B)** Scatterplot showing the correlation of fitness scores for individual mutations between N1/Mar43 and N1/Geo06. Pearson correlation coefficient (R) is indicated. Mutations of interested are indicated. **(C and D)** The replication fitness effects of mutations at **(C)** H412a and **(D)** P459 residue across different human H1N1 strains were analyzed by virus rescue experiment. Mutation was individually introduced into NA of indicated H1N1 strains. Viral titer was measured by TCID_50_ assay. Each bar represents the mean of three independent biological replicates. **(E and H)** Interactions between neighboring residues with **(E)** H412a and **(H)** P459 is shown, respectively. **(F and G)** Sequence logos showing the sequence conservation for residues surrounding **(F)** 412a and **(G)** 459, respectively. NA sequences from N1/PR8 and 11 human H1N1 strains indicated in **(D)** were analyzed. **(I)** Phylogenetic tree of 9 human H1N1 NAs, with their NA activity and thermostability shown on the right as heatmaps. NA activity was quantitated by ELLA assay (red gradient scale). NA thermostability was measured by thermal shift assay (blue gradient scale). Of note, N1/Mar43 and A/Bel/1942 are excluded because their thermal unfolding profiles exhibited two distinct transitions.

### Mutational fitness landscape of H1N1 NA is influenced by epistasis

To further identify mutations involving in epistasis, we compared the mutational fitness landscapes between N1/Mar43 and N1/Geo06 NAs and identified mutations that were beneficial in one strain but deleterious in the other. Specifically, P459N/E/D mutations were beneficial in N1/Mar43 but detrimental in N1/Geo06, whereas H412aM was beneficial in N1/Geo06 but deleterious in N1/Mar43 (**Figure 3B**). These reciprocal effects highlighted the role of epistasis in shaping strain-dependent evolutionary constraints.

To determine whether the differential mutational fitness effects at H412a and P459 were unique to N1/Geo06 or represented a broader evolutionary feature of H1N1 NAs, mutagenesis was performed across a panel of human H1N1 strains from 1942 to 1999. Our virus rescue experiment showed that H412aM abolished the viral replication for all tested strains except N1/Geo06 (**Figure 3C**), whereas the replication fitness effects of P459D/N had a smaller magnitude in general, yet remained strain-dependent (**Figure 3D**). To substantiate these findings, we performed deep mutational scanning at both positions in seven human H1N1 strains. Consistent with the virus rescue data, residue H412a was highly intolerant to mutations in these seven strains, whereas the strain-dependent replication fitness effects of mutations at P459 were apparent (**Figure S6A-B**). Structural analysis showed that H412a does not form direct local interactions and that surrounding residues are highly conserved (**Figure 3E-F**). Similarly, although structural modelling of P459D/N/E showed the introduction of interprotomer H-bonds, its surround residues are also highly conserved (**Figure 3G-H and S6C-F**). These observations suggested that the strain-dependent fitness effects of H412aM and P459D/N/E arose from long-range epistasis.

We next examined whether NA protein stability and enzymatic activity could help explain the differential fitness effects of H412aM and P459D/N across strains. Recombinant NA proteins from different H1N1 strains exhibited differences in thermostability and enzymatic activity, as determined by thermal shift assay and enzyme-linked lectin (ELLA) assay, respectively (**Figure 3I**). While thermostability and enzymatic activity moderately correlated (ρ = 0.52) (**Figure S6G**), neither protein stability nor enzymatic activity explained the strain-dependent fitness effects of H412aM or P459D/N (**Figure S6H-I**). For example, N1/Geo06 did not exhibit exceptional thermostability or enzymatic activity despite uniquely tolerating H412aM (**Figure 3C and Figure S6H**). Similarly, N1/NC99 showed biochemical properties comparable to other strains despite uniquely failing to tolerate P459D (**Figure 3D**). Collectively, these results showed that differences in the NA thermostability and enzymatic activity were insufficient to explain the strain-dependent fitness effects of H412aM and P459D/N.

### Distinct distributions of fitness effects across N1 and N2 strains

To understand how the evolutionary potential of NA varies among strains, we compared the distributions of fitness effects of the three N1 NAs analyzed in the current study (N1/PR8, N1/Mar43, and N1/Geo06) and that of H3N2 A/Moscow/10/1999 (N2/Mos99) NA from our previous study^15^. We observed that the distribution of fitness effects of missense mutations differed substantially among strains (**Figure 4A**). While the human H1N1 isolates N1/Mar43 and N1/Geo06 showed a unimodal distribution centered near 0, N1/PR8 and N2/Mos99 exhibited a bimodal distribution with distinct peaks at around 0 and 1. This bimodality reflected a higher proportion of well-tolerated mutations in the latter strains. For example, compared to N2/Mos99 NA, N1/Mar43 NA showed lower mutational tolerance at antigenic, exposed and buried residues (**Figure 4B and Figure S7A**). Additionally, the NA mutational fitness effects only moderately correlated between N2/Mos99 and different N1 NAs (R = 0.35 to 0.49) (**Figure 4C and Figure S7B-C**). These correlation coefficients were similar to those observed among the N1 strains themselves (**Figure 3B and Figure S5E-F**), indicating that the difference in NA mutational fitness landscapes between distinct strains within a subtype could be as large as that between strains from different subtypes.

**Figure 4.**
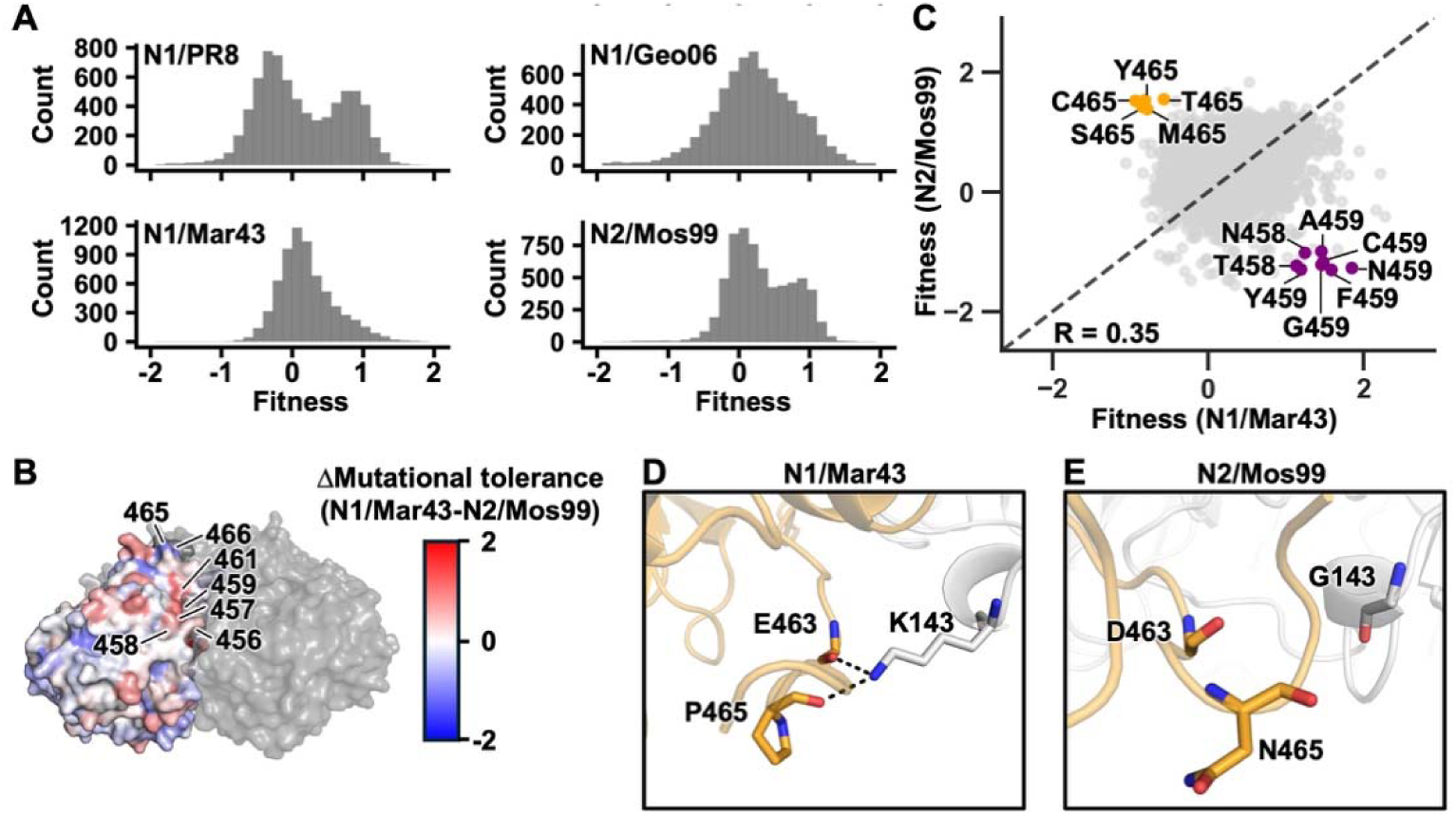
Comparison of NA mutational effects between human H1N1 and H3N2 viruses. **(A)** Histograms showing the distributions of fitness scores for missense mutations of the indicated strains. **(B)** Difference of mutational tolerance between N1/Mar43 and N2/Mos99 mapped onto the cryo-EM structure of N1/Mar43 NA. One protomer is displayed in surface representation with residues colored according to mutational tolerance, while the remaining three protomers are shown in gray. **(C)** Scatterplot showing the correlation of fitness scores for individual mutations between N1/Mar43 and N2/Mos99. Pearson correlation coefficient (R) is indicated. Mutations of interested are indicated. **(D)** Interactions between P465 and neighboring residues in N1/Mar43 NA is shown. **(E)** Interactions between N465 and neighboring residues in N2/Mos99 NA (PDB: 7U4F)^17^ is shown.

The major differences in the mutational fitness landscapes between N1 NAs and N2/Mos99 NA were at the C-terminus (**Figure 4B-C and S7B-C**). Residue 465 in N2/Mos99 was more permissive to mutations than N1 NAs. Structure analysis showed that P465 formed an interprotomer H-bond with K143 in N1/Mar43 NA (**Figure 4D**), whereas N465 in N2/Mos99 NA lacked direct local interactions (**Figure 4E**). To determine whether these features were conserved, we further analyzed available human N1 and N2 NA crystal structures. Most N1 structures retained the interprotomer interaction observed in N1/Mar43 P465 (**Figure S7D)**, whereas all N2 NA structures analyzed formed intraprotomer interactions at this position (**Figure S7F)**. While this finding illustrated that the absence of local interactions in N2/Mos99 was a strain-specific structural feature, it also suggested that the epistatic effects at the C-terminal residues between N1 and N2 NAs were due to differences in the interprotomer interactions.

## DISCUSSION

Both HA and NA are major antigens of IAV. However, while the mutational fitness landscape of HA has been extensively characterized by deep mutational scanning^19–22^, that of NA is much less well characterized, despite it being an important target for influenza countermeasures. In this study, we measured the replication fitness effects of nearly all possible amino acid mutations across the head domains of three different H1N1 NAs. Our results revealed that while the evolutionary constraints of many residues were similar across strains, the mutational tolerance of some residues was strongly influenced by epistasis. We further analyzed the biophysical determinants of mutational fitness and epistasis. Overall, our study not only provides mechanistic insights into the evolutionary constraints of NA, but also offers a dataset that is valuable to the field, especially since increasing attention has focused on NA as a target for influenza vaccine development.

One important finding is that the strong concordance between NA mutational fitness landscapes *in vitro* and *in vivo*. A similar observation was previously reported in a deep mutational scanning study of H1N1 A/WSN/1933 NA using a mutant library generated by error-prone PCR^23^. These observations suggest that the functional and biophysical constraints of NA evolution in a commonly used cell culture model (MDCK-SIAT1 cells) and in host respiratory tract are similar. Nevertheless, replication fitness is not equivalent to transmission fitness. Viral replication in the host respiratory tract is influenced by immune responses and host-specific physiological conditions^24–27^, whereas transmission also depends on factors such as viral shedding, stability in respiratory secretions, and environmental persistence^28–32^. Thus, although mouse model is typically considered as a physiologically relevant measure of viral replication, additional selective pressures associated with transmission may further shape the evolution of NA in circulating strains. However, the relationship between replication fitness and transmission fitness is unclear and warrants further analysis.

Another notable finding from our study is the substantial difference in the distribution of fitness effects across N1 and N2 strains. In particular, the bimodal fitness distributions observed in N1/PR8 and N2/Mos99 suggest that certain NAs possess greater mutational tolerance and therefore have higher evolutionary potential. Our previous studies have shown that the mutational tolerance of HA is influenced by NA activity^33^, due to the functional balance between HA and NA^3^. In our present study, HA from PR8 or A/WSN/33 was used for all deep mutational scanning experiments. It is possible that the mutational tolerance of N1/Mar43 and N1/Geo06 would alter when paired with their corresponding native HAs. At the same time, mutational tolerance is known to be influenced by protein thermostability^34^. Because the mutational tolerance of NA underpins the potential for acquiring antibody escape and drug resistance, future studies should investigate its underlying determinants.

A number of heterosubtypic antibodies to NA have been identified in recent years^35–42^, highlighting the possibility of leveraging NA for the development of broadly protective influenza vaccines^6^. However, the escape mutation profiles for these NA antibodies remain unclear. Previous studies have shown the feasibility of using deep mutational scanning to identify escape mutations against broadly neutralizing HA antibodies^43,44^. We postulate that the NA mutant libraries constructed in this study can similarly be used to identify escape mutations against NA antibodies, which will have important implications for the development of NA-based broadly protective vaccines. We anticipate that future deep mutational scanning studies of NA will continue to advance our understanding of its evolution and its role as a vaccine target.

## ACKNOWLEDGEMENTS

We thank the Roy J. Carver Biotechnology Center at the University of Illinois Urbana-Champaign for assistance with next-generation sequencing. We gratefully acknowledge the use of Case Western Reserve University Cryo-Electron Microscopy Facility (supported by NIH grant S10OD032437) and the High-Performance Computing Resource in the Core Facility for Advanced Research Computing. This project has been funded in part with Federal funds from the National Institute of Allergy and Infectious Diseases, National Institutes of Health, Department of Health and Human Services, under Contract No. 75N93021C00015 (N.C.W.), as well as with the Vallee Scholars Program (N.C.W.) and the Carl R. Woese Institute for Genomic Biology Postdoctoral Fellowship (H.L.).

## AUTHOR CONTRIBUTIONS

Q.W.T., H.L., and N.C.W. conceived and designed the study. Q.W.T., constructed the NA deep mutational scanning library and prepared the sequencing libraries. Q.W.T., W.L., and N.C.W. performed data analysis. H.L. performed mice experiments and recombinant protein expressions. Z.M., R.L., and X.D. solved the cryo-EM structure of N1/Mar43 NA. Q.W.T., H.L., K.J.M., Y.W.H., J.J.H., A.B.G., A.M., and E.X.M. performed other experiments. Q.W.T., H.L., and N.C.W. wrote the paper and all authors reviewed and/or edited the paper.

## METHODS

### Cells cultures

Madin-Darby canine kidney cells with stable expression of human α-2,6-sialyltransferase (MDCK-SIAT1) (Sigma, catalog #: 05071502-1VL) and human embryonic kidney HEK293T (ATCC, catalog #: CRL-3216) cells were maintained in Dulbecco’s modified Eagle’s medium (DMEM; Gibco) supplemented with 10% fetal bovine serum (FBS; Gibco), 100 U/mL Penicillin-Streptomycin. Cells were cultured in humidified incubators at 37°C with 5% CO_2_. SF9 cells (Spodoptera frugiperda ovarian cells, female, ATCC) were maintained in Sf-900 II SFM medium (Thermo Fisher Scientific).

### Mutant library construction

The mutant library was constructed based on the pHW2000 eight-plasmid reverse genetics system for influenza virus^45^. N1/PR8 NA-encoding pHW2000 plasmid (pHW2000-PR8-NA), N1/Mar43 NA-encoding pHW2000 plasmid (pHW2000-Marton43-NA) or N1/Geo06 NA-encoding pHW2000 plasmid (pHW2000-Geo06-NA) was used as a PCR template to generate a linearized vector and a library of mutant inserts. The linearized vector was generated using 5’-CGT ACG TCT CAA CAC ACG GAG CGC CCG GGG CCC TCT-3’ and 5’-CGT ACG TCT CAT AAC TGC TAG CGT TAA CAG CTT GGG-3’ as primers. For insert generation, two batches of PCRs were performed, followed by overlapping PCRs. The first batch of PCRs consisted of 48 reactions, each with an equal molar mix of eight forward primers, as the forward primer and a universal reverse primer 5’-CGT ACG TCT CAG TGT GGC TGC GAT GGT GGC GTT T-3’. The forward primers for the first batch of PCRs are listed in **Table S3**. These forward primers were named as cassetteX_N, in which X represents the cassette ID and N represents the primer number. Forward primers with the same cassette ID were mixed at equal molar ratios and used in the same single PCR. The second batch of PCRs consisted of another 48 reactions, each with a universal forward primer 5’-CGT ACG TCT CAG TTA ACC GGA GTA CTG GTC GAC C-3’. and unique reverse primers as listed in **Table S3**. Subsequently, 48 overlapping PCRs were performed using the universal forward primer and reverse primer. For each overlapping PCR, the template was a mixture of 10 ng each of the corresponding products from the first and second batches of PCRs. The complete insert was an equal molar mix of the products of these 48 overlapping PCRs. All PCRs were performed using PrimeSTAR Max polymerase (Takara Bio) according to the manufacturer’s instructions. PCR products were purified using Monarch DNA Gel Extraction Kit (New England Biolabs). Both the vector and the complete insert were digested by BsmBI (New England Biolabs) and ligated using T4 DNA ligase (New England Biolabs). The ligated product was transformed into MegaX DH10B T1R cells (Thermo Fisher Scientific). At least one million colonies were collected. Plasmid mutant libraries were purified from the bacteria colonies using PureLink HiPure Expi Plasmid Purification Kit (Thermo Fisher Scientific).

For N1/PR8, the same general strategy was used with different vector and insert primers and different cloning method. The linearized vector was generated using 5’-CGT ACG TCT CAA CAC ACG GAG CGC CCG GGG CCC TCT-3’ and 5’-CGT ACG TCT CAC TAG AGG ATC CCC GGG TAC CGA GCT-3’ as primers. The universal forward and reverse primers 5’-CGT ACG TCT CAC TCT AGA CCG GAG TAC TGG TCG ACC-3’ and 5’-CGT ACG TCT CAG TGT GGC TGC GAT GGT GGC GTT T-3’, respectively. Both the vector and the complete insert were ligated using NEBuilder HiFi DNA assembly master mix (New England Biolabs).

### Rescuing and passaging the mutant library

The virus mutant library was rescued by transfecting a co-culture of HEK293T and MDCK-SIAT1 cells (ratio of 6:1) at 60% confluence in a T175 flask (175 cm^2^) with seven internal segments (PB2, PB1, PA, NP, HA, M, and NS) from H1N1 A/Puerto Rico/8/1934 for N1/PR8 libraries, A/WSN/33 for N1/Mar43 and N1/Geo06 libraries. Transfection was performed using Lipofectamine 2000 (Thermo Fisher Scientific) according to the manufacturer’s instructions. At 24 h post-transfection, cells were washed twice with PBS and cell culture medium was replaced with OPTI-MEM medium supplemented with 0.8 μg/mL TPCK-trypsin. The virus mutant library was harvested at 72 h post-transfection, titered by TCID_50_ assay using MDCK-SIAT1 cells, and stored at –80°C until use. To passage the virus mutant libraries, MDCK-SIAT1 cells in a T175 flask were washed twice with PBS and then infected with a multiplicity of infection (MOI) of 0.02 in OPTI-MEM medium supplemented with 0.8 μg/mL TPCK-trypsin. At 2 h post-infection, infected cells were washed twice with PBS and fresh OPTI-MEM medium supplemented with 0.8 μg/mL TPCK-trypsin was added to the cells. At 24 h post-infection, supernatant was harvested. Each replicate was transfected and passaged independently. For N1/PR8 mutant library in mice, the N1/PR8 P1 virus libraries were further passaged in mice via intranasal infection with 100 TCID₅₀, with lung tissues harvested on day 3 post-infection. For each biological replicate, lungs from five mice were pooled, and two independent replicates were processed in parallel.

### Sequencing library preparation

Viral RNA of the post-passaged mutant library was extracted using QIAamp Viral RNA Mini Kit (QIAGEN). The extracted RNA was then reverse transcribed to cDNA using Superscript III reverse transcriptase (Thermo Fisher Scientific). The wild-type (WT) plasmid, plasmid mutant library, and the cDNA from the post-infection viral mutant library were amplified by PCR to add part of the adapter sequence required for Illumina sequencing. Six amplicon PCRs were performed for each sample using the primers listed in **Table S3**. A total of 12 Ns were included in each amplicon product as unique molecular identifiers (UMIs) to distinguish true mutations from sequencing errors^46–48^. For each sample, the six amplicon PCR products were mixed at equal molar ratio. Subsequently, 1 million copies of mixed amplicon PCR products were used as template for a second PCR to add the rest of the adapter sequence and index to the amplicon using primers: 5’-AAT GAT ACG GCG ACC ACC GAG ATC TAC ACX XXX XXX XAC ACT CTT TCC CTA CAC GAC GCT-3’, and 5’-CAA GCA GAA GAC GGC ATA CGA GAT XXX XXX XXG TGA CTG GAG TTC AGA CGT GTG CT-3’. Positions annotated by an “X” represented the nucleotides for the index sequence. All PCRs were performed using KOD Hot Start DNA polymerase (Millipore Sigma) according to the manufacturer’s instructions. PCR products were purified by Monarch DNA Gel Extraction Kit (New England Biolabs). The final PCR products were submitted for next-generation sequencing using one lane of NovaSeq SP PE150.

### Sequencing data analysis

Sequencing data was obtained in FASTQ format and analyzed using a custom snakemake pipeline^49^. Firstly, forward and reverse consensus reads were merged by PEAR using parameters: -v 20 -j 6. Paired-end reads with the same UMI were then grouped together. Groups with only two or fewer reads were discarded. Groups with reads that had different lengths were also discarded. A consensus read was generated for each group of reads with an 80% minimum frequency cutoff. In other words, for each position, the same nucleotide needed to be shared among at least 80% of the reads, otherwise the group would be discarded. Subsequently, each consensus read was assigned to one of the six amplicons based on the primer sequence. Consensus reads that were not assigned to any amplicon were discarded. Based on the mutagenic primers and reference sequence, mutant reference sequences were generated iteratively using a Python script to construct a mutation library. Merged reads were then aligned to this library using BWA^50^, and only perfect matches were retained to quantify each mutation type. The frequency of each mutation in individual samples was computed. The frequency of mutation *i* in sample *s* was calculated as follows:

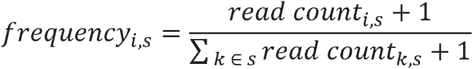

A mutation would be discarded if its frequency in the plasmid mutant library was not at least 6-fold higher than its frequency in the WT sample. This filter aimed to prevent error hotspots in reverse transcription, PCR, and next-generation sequencing from confounding our fitness calculation. In addition, a mutation would be discarded if its read count in the plasmid mutant library was 10 or less. After these filter steps, fitness calculation was performed. For each replicate, the enrichment and fitness scores of a mutation *i* in amplicon *amp* was calculated as follows:

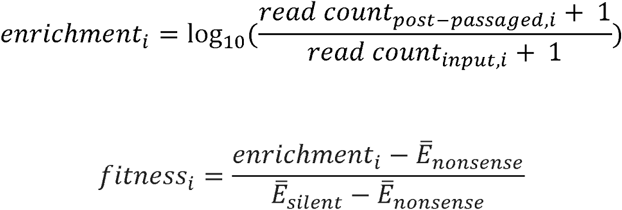

where the *read count_input_*_;*i*_ represents the read count of mutation *i* in the plasmid mutant library, *read count_post-passaged_*_;_*i* represents the read count of mutation *i* in the post-passaged mutant library *E̅_nonsense_*, represents the mean enrichment score of nonsense mutations and *E̅_silent_*represents the mean enrichment score of silent mutations. A pseudocount of 1 was added to the read counts to avoid division by zero. The fitness score for mutation *i* was scaled using nonsense and silent mutations, such that the mean fitness score for nonsense mutations was 0 and the mean fitness score for silent mutations was 1. The final fitness score was computed as the averaged fitness from the two biological replicates.

The mutational tolerance for each residue was quantified based on the average fitness scores of all mutations at a given residue, and was calculated as follows:

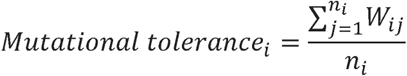

where the *W_ij_* represents the fitness of mutation *j* at residue *i*, and *n_i_* represents the number of mutations measured at residue *i*.

### Rescue of NA mutant viruses

N1/PR8 mutant viruses were generated using the pHW2000 eight-plasmid reverse genetic system. Briefly, the virus mutant library was rescued by transfecting a co-culture of HEK293T and MDCK-SIAT1 cells (ratio of 6:1) at 60% confluence in a T175 flask (175 cm^2^) with seven internal segments (PB2, PB1, PA, NP, HA, M, and NS) from PR8. Transfection was performed using Lipofectamine 2000 (Thermo Fisher Scientific) according to the manufacturer’s instructions. At 24 h post-transfection, cells were washed twice with PBS and cell culture medium was replaced with OPTI-MEM medium supplemented with 0.8 μg/mL TPCK-trypsin. The virus mutant library was harvested at 72 h post-transfection, titered by TCID_50_ assay using MDCK-SIAT1 cells.

### Flow cytometry analysis of NA expression

4×10^5^ HEK293T cells were seeded in 6-well plates and grown overnight at 37°C and 5% CO_2_ in a humidified incubator. Cells were transfected with 2 μg of pCAGGS plasmids encoding the C-terminal Myc-tagged NA from the strain of interest using Lipofectamine 2000 (Thermo Fisher Scientific) according to the manufacturer’s instructions. At 24 h post-transfection, medium was discarded, and cells were washed once with PBS. Cells were subsequently detached with versene (Thermo Fisher Scientific) and resuspended in fluorescence-activated single cell sorting (FACS) buffer (50 mM EDTA, 2% v/v FBS in DMEM with high glucose and HEPES, without phenol red). Cells were then pelleted via centrifugation at 300 ×*g* for 5 min at 4°C and the supernatant was aspirated. In the subsequent steps, cells and all reagents were kept on ice.

For surface staining, cell pellets were resuspended in 1 mL FACS buffer. Cells were incubated with 1 μg/mL APC anti-c-Myc (BioLegend, catalog #: 926810) at 4°C with rocking for 1 h. Subsequently, cells were pelleted via centrifugation at 300 ×*g* for 5 min at 4°C and the supernatant was aspirated. Cells were washed with 1 mL FACS buffer, pelleted, and resuspended in 1 mL FACS buffer before flow cytometry analysis.

For intracellular staining, cell pellets were fixed in 300 μL of 1% v/v paraformaldehyde (in PBS) and incubated at 4°C with rocking for 30 min. Cells were pelleted, washed with 1 mL FACS buffer, permeabilized in 300 μL of 0.1% v/v Triton X-100 (in PBS) and incubated at 4°C with rocking for 30 min. Cells were pelleted, washed with 1 mL FACS buffer and incubated with 1 μg/mL APC anti-c-Myc (BioLegend, catalog #: 926810) at 4°C with rocking for 1 h. Then, cells were pelleted, washed with 1 mL FACS buffer, and resuspended in 1 mL FACS buffer before flow cytometry analysis. For flow cytometry analysis, cells were analyzed using NovoCyte Opteon spectral flow cytometer (Agilent). Data were analyzed using FlowJo v10.8 software (BD Life Sciences).

### ER stress analysis

ER stress induction was measured using XBP1 mRNA splicing luciferase reporter^18,51^. Briefly, HEK293T cells were seeded in 24-well plates and grown overnight at 37°C and 5% CO_2_ in a humidified incubator. Cells were then transfected with the reporter plasmid and NA expression plasmids. The cells were lysed at 24 h post transfection, and the luciferase assay was performed using a dual-luciferase reporter assay system (Promega).

### Protein expression and purification

The NA head domains were fused to an N-terminal gp67 signal peptide, 6× His-tag, a vasodilator-stimulated phosphoprotein (VASP) tetramerization domain, and a thrombin cleavage site. Recombinant bacmid DNA that carried NA ectodomain from the strain of interest was generated using the Bac-to-Bac system (Thermo Fisher Scientific) according to the manufacturer’s instructions. Baculovirus was generated by transfecting the purified bacmid DNA into adherent Sf9 cells using Cellfectin reagent (Thermo Fisher Scientific) according to the manufacturer’s instructions. The baculovirus was further amplified by passaging in adherent Sf9 cells at a multiplicity of infection (MOI) of 1. Recombinant NA head domains were expressed by infecting 1 L of suspension Sf9 cells at an MOI of 1. On day 3 post-infection, Sf9 cells were pelleted by centrifugation at 4,000 ×*g* for 25 min, and soluble recombinant NA was purified from the supernatant by affinity chromatography using Ni Sepharose excel resin (Cytiva) and then size exclusion chromatography using a HiLoad 16/100 Superdex 200 prep grade column (Cytiva) in 20 mM Tris-HCl pH 8.0, 100 mM NaCl, and 10 mM CaCl_2_.

### Enzyme-linked lectin assay (ELLA)

ELLA experiments were performed described^52^. Briefly, each well of a 96-well microtiter plate (Thermo Fisher) was coated with 100 μL fetuin (Sigma) at 25 mg/mL in coating buffer (KPL coating solution; SeraCare) at 4°C overnight. The next day, 50 μL antibodies at the indicated concentrations in 2-(N-morpholino) ethanesulfonic acid (MES) at pH 6.5, 20 mM CaCl_2_, 1% bovine serum albumin and 0.5% Tween 20 were mixed an equal volume of NA (0.032 μg/well). This mixture was added to the fetuin-coated plate and incubated for 18 h at 37°C. The plate was then washed six times with PBS with 0.05% Tween 20. Subsequently, 100 μL of HRP-conjugated peanut agglutinin lectin (PNA-HRPO, Sigma–Aldrich) in MES at pH 6.5 with CaCl_2_ and 1% bovine serum albumin was added and incubated for 2 h at room temperature in the dark. The plate was washed six times and developed with 3,3’5,5’-tetrame-thylbenzidine (TMB) ELISA substrate (Sigma). Absorbance was read at 450 nm using a BioTek synergy HTX multimode reader (Agilent). Data points were analyzed using Prism software and the 50% inhibition concentration (IC_50_) was defined as the concentration at which 50% of the NA activity was inhibited compared to the negative control.

### Thermal shift assay

5 µg protein was mixed with 5× SYPRO orange (Thermo Fisher Scientific) in 20 mM Tris-HCl pH 8.0, 100 mM NaCl, and 10 mM CaCl_2_ at a final volume of 25 µL. The sample mixture was then transferred into an optically-clear 96-well PCR plate (Bio-Rad). SYPRO orange fluorescence data in relative fluorescence unit (RFU) was collected from 10°C to 95°C using a CFX Connect Real-Time PCR Detection System (Bio-Rad). The temperature corresponding to the lowest point of the first derivative −d(RFU)/dT was determined to be the T_m_.

### Cryo-EM sample preparation, data collection, and data processing

The peak fractions of the NA from size exclusion chromatography were concentrated to around 1 mg/mL for cryo-EM sample preparation. Cryo-EM grids were prepared using a Vitrobot Mark IV machine. An aliquot of 3.5 μL sample was applied to a 300-mesh Quantifoil R1.2/1.3 Cu grid pre-treated with glow-discharge. Excess liquid was blotted away using filter paper with blotting force −5 and blotting time 3 s. The grid was plunge frozen in liquid ethane. Data collection was done with serialEM^53^ on a Titan Krios microscope equipped with Gatan BioQuantum K3 imaging filter and camera. A 20 eV slit was used for the filter. Images were recorded at 130,000x magnification, corresponding to a pixel size of 0.33 Å/pix at super-resolution mode of the camera. A defocus range of −0.8 μm to −1.5 μm was set. A total dose of 50 e-/Å^2^ of each exposure was fractionated into 50 frames. The first two frames of the movie stack were excluded in motion-correction. Cryo-EM data processing was performed on the fly with cryoSPARC Live^54^, following regular single-particle procedures. The model of NA (PDB: 9DPC)^55^ was fitted into the cryo-EM density map using UCSF Chimera^56^. The model was manually adjusted in Coot^57^ and refined with Phenix real-space refinement program^58^. This process was iterated for several cycles until no significant improvement of the model was observed.

## Data availability

Raw sequencing data have been submitted to the NIH Short Read Archive under accession numbers: BioProject PRJNA1519971. The cryo-EM density map and coordinate of N1/Mar43 NA have been deposited to the Electron Microscopy Data Bank (EMDB) with accession code EMD-75452 and the Protein Data Bank (PDB) with accession code 10TB, respectively.

## Code availability

Custom python scripts for all analyses have been deposited to: https://github.com/Karobben/DMS-H1N1-NA

---

**Figure S1.**
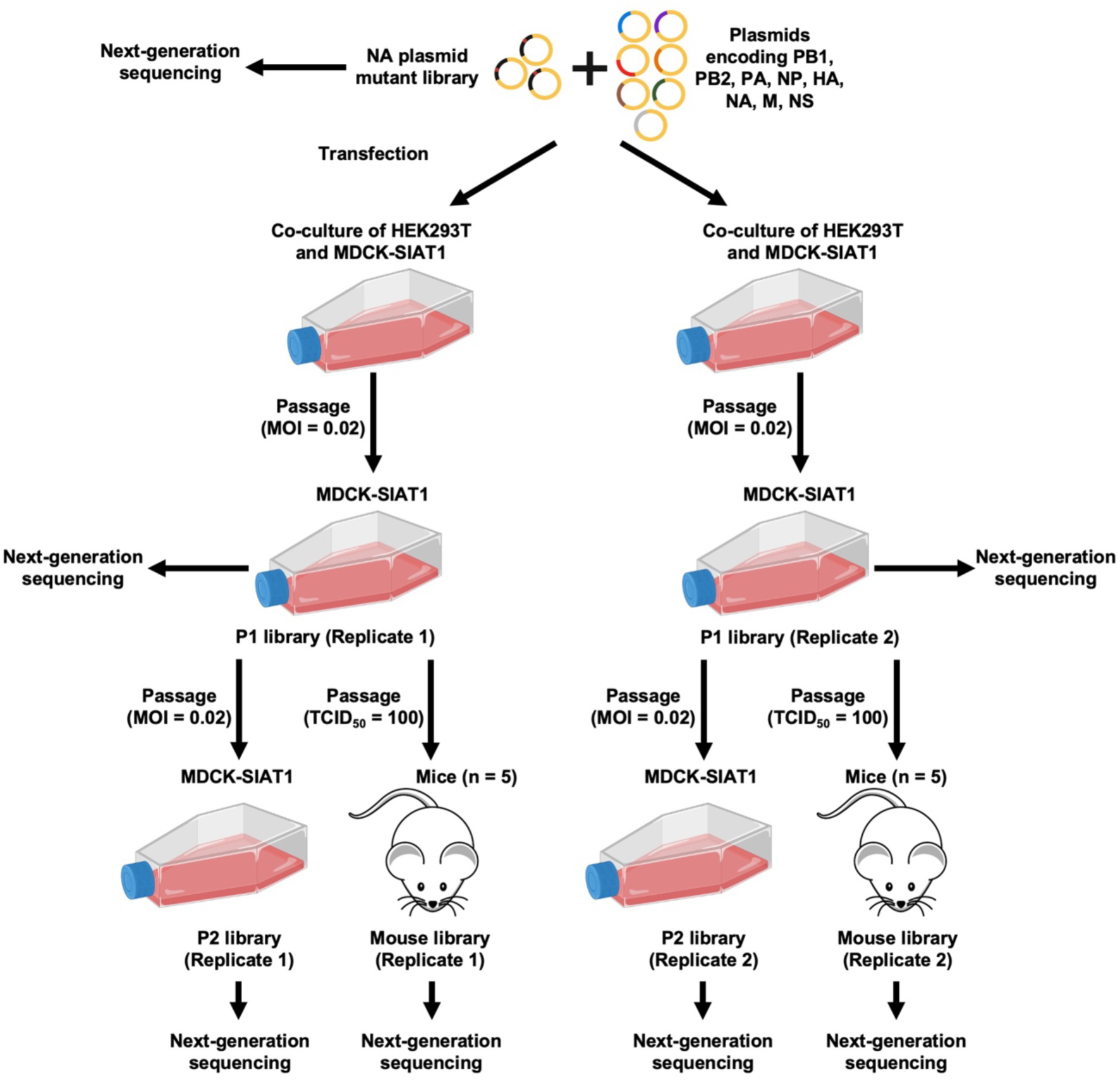
Experimental design. The PR8 NA virus mutant library was rescued by transfecting the PR8 NA plasmid mutant library along with plasmids encoding seven internal segments from A/Puerto Rico/8/1934 (H1N1) into a co-culture of HEK293T cells and MDCK-SIAT1 cells. The PR8 NA virus mutant library was then passaged in MDCK-SIAT1 cells at a multiplicity of infection (MOI) of 0.02 to generate the post-passaged P1 mutant library. The P1 library was further passaged either in MDCK-SIAT1 cells at MOI 0.02 (P2 library) or in mice at 100 TCID_50_ (mouse library). A biological replicate of transfection and passaging was performed using the same NA plasmid mutant library. For each mouse biological replicate, lungs from five mice were pooled. The NA plasmid mutant library and both biological replicates of the P1, P2, and mouse libraries were analyzed by next-generation sequencing.

**Figure S2.**
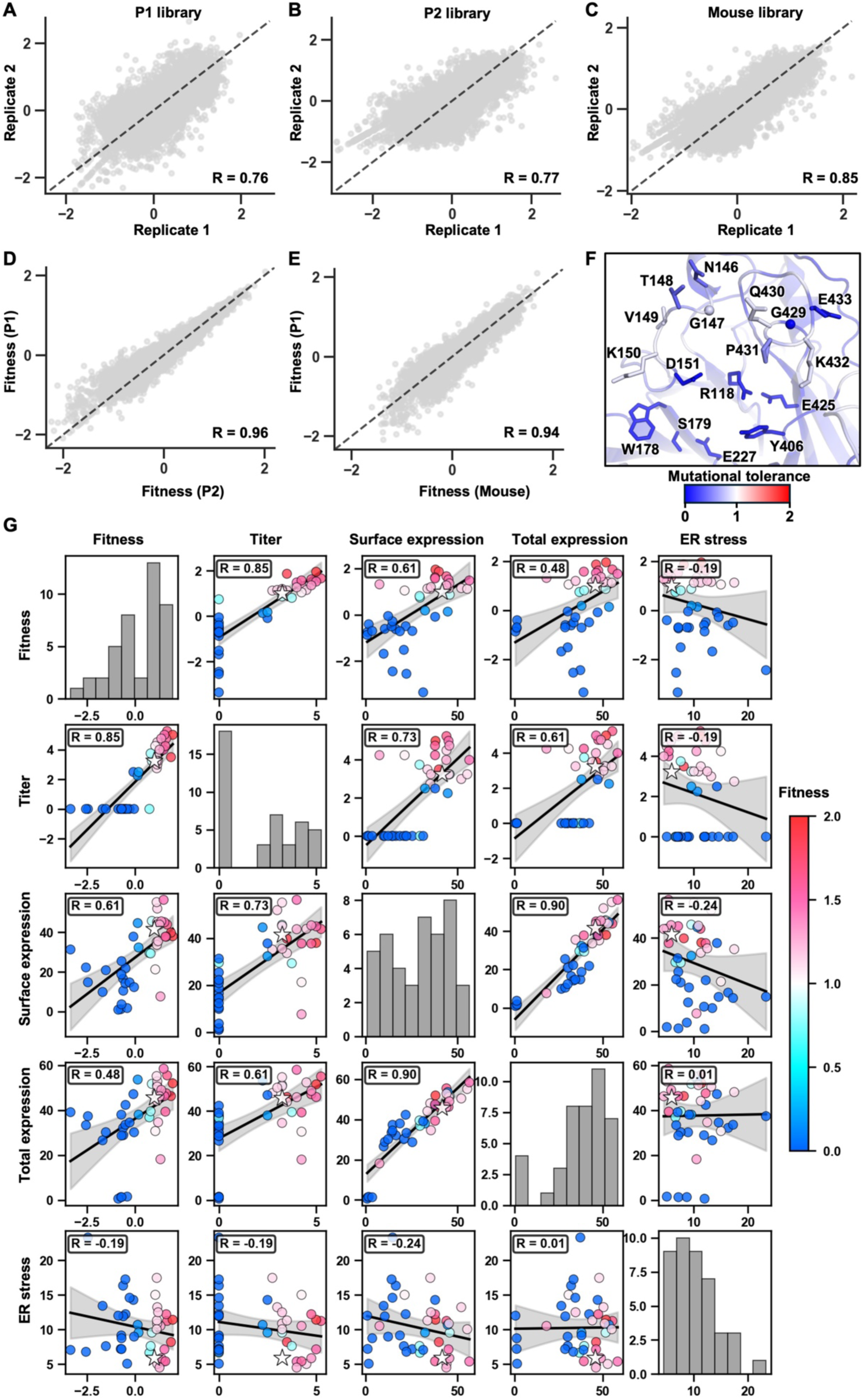
Reproducibility and quality of N1/PR8 deep mutational scanning data. **(A-C)** Correlation of fitness scores for individual mutations between two biological replicates for **(A)** P1 library, **(B)** P2 library, and **(C)** mouse library is shown as a scatterplot. **(D and E)** Scatterplot showing the correlation of fitness scores for individual mutations between P1 library and **(D)** P2 library, or **(E)** mouse library. **(F)** Mutational tolerance across NA residues of N1/PR8 P1 library mapped onto the NA structure from H1N1 A/Brevig Misson/1/1918 (PDB: 8E6K)^1^. Catalytic residues (R118 and Y406), framework residues (W178, S179, E277, and E425), 150 loop (G147 and K150) and 430-loops (Q430 and K432) are labeled. **(G)** Scatterplots showing the relationships among fitness score, viral titer, NA surface expression, NA total expression, and ER stress. Individual data points are color mapped, where red and blue indicate beneficial and deleterious mutations, respectively. WT is represented by a star. **(A-E and G)** The Pearson correlation coefficient (R) is indicated.

**Figure S3.**
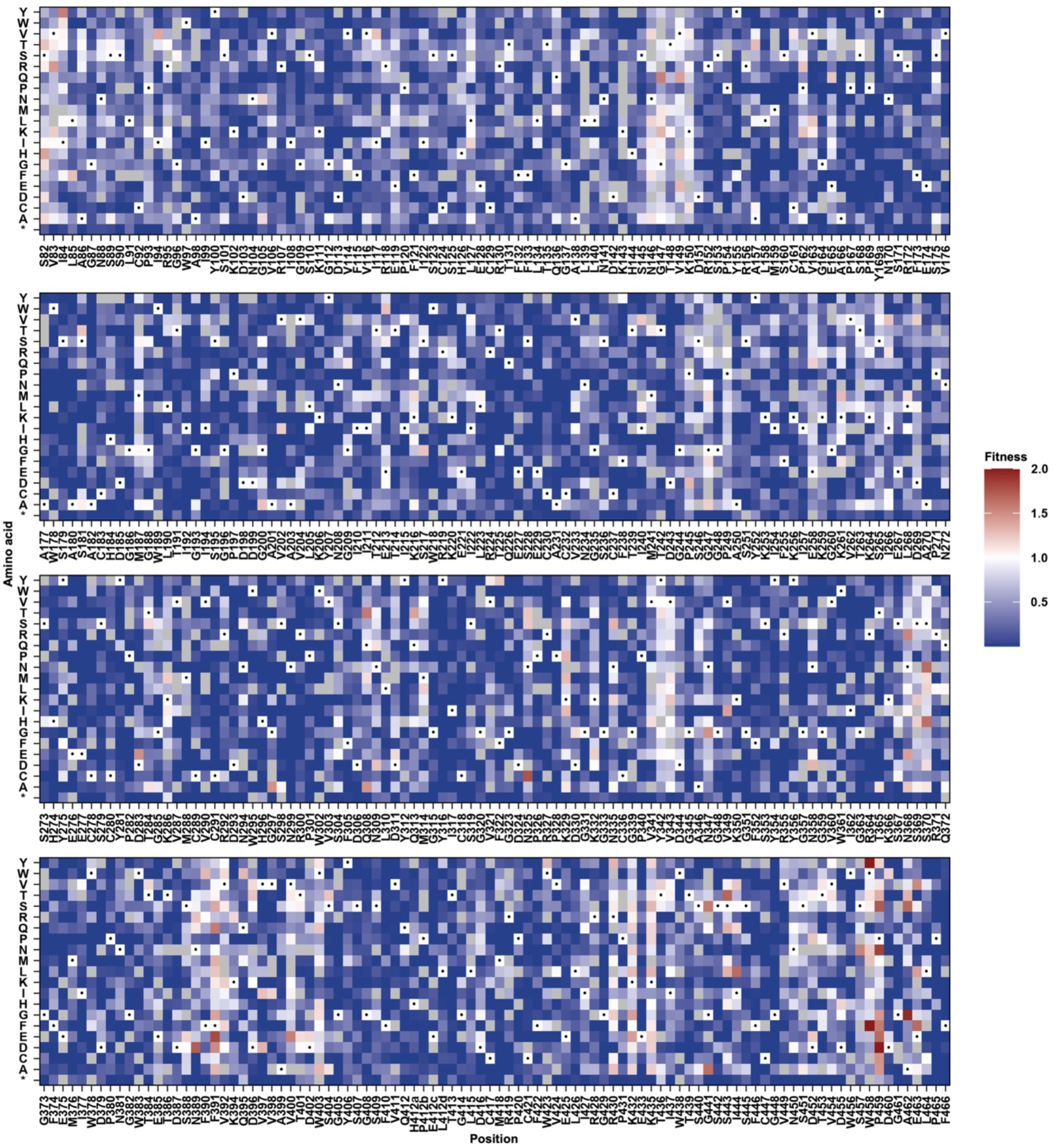
Deep mutational scanning of N1/Mar43 NA. The replication fitness effects of individual mutations in the N1/Mar43 NA library were measured by deep mutational scanning and fitness scores are shown as a heatmap. Wild-type (WT) amino acids are indicated by a black circle. Mutations in gray were excluded in our data analysis due to low input count. Stop codon is annotated as *.

**Figure S4.**
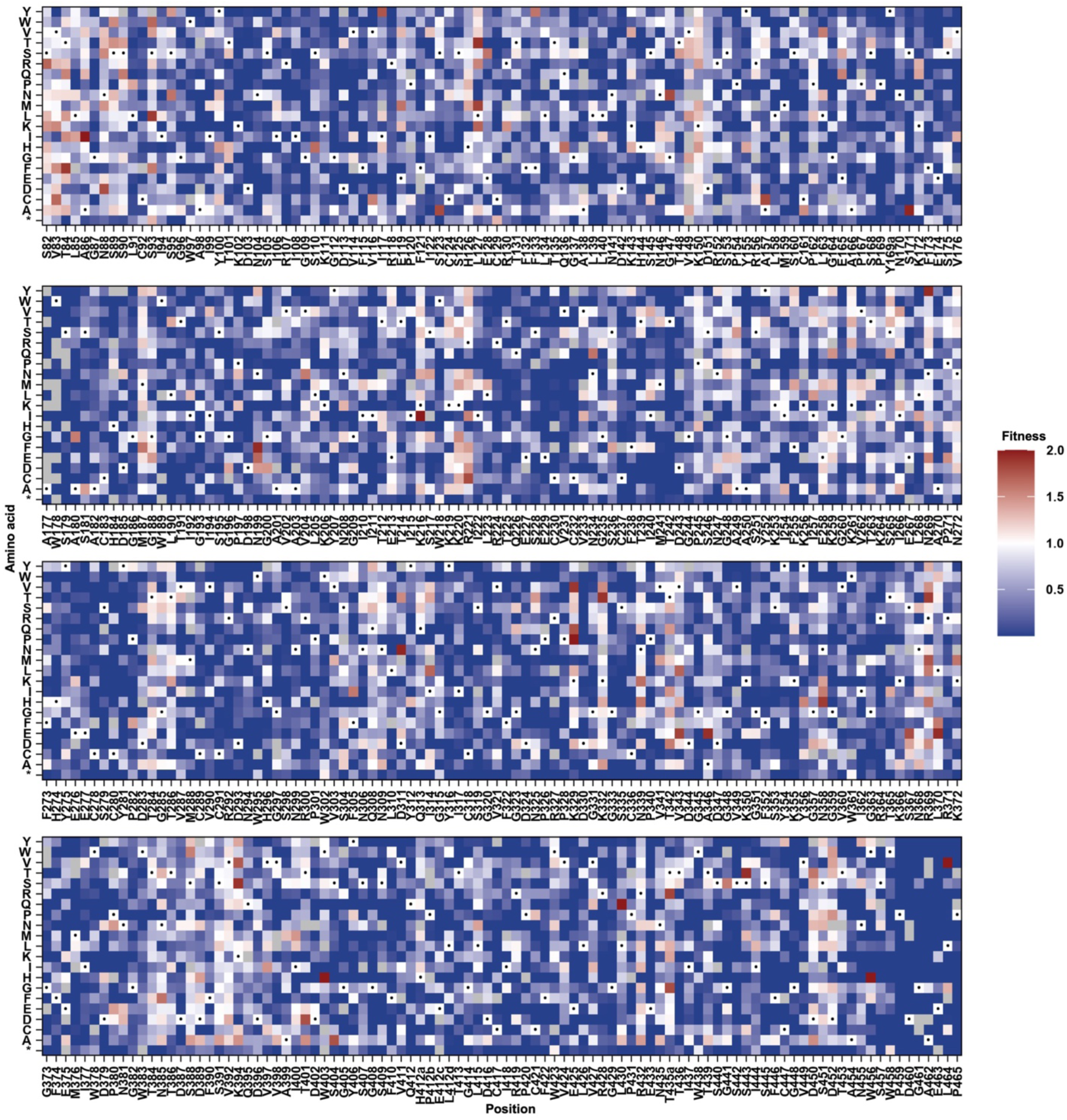
Deep mutational scanning of N1/Geo06 NA. The replication fitness effects of individual mutations in the N1/Geo06 NA library were measured by deep mutational scanning and fitness scores are shown as a heatmap. WT amino acids are indicated by a black circle. Mutations in gray were excluded in our data analysis due to low input count. Stop codon is annotated as *.

**Figure S5.**
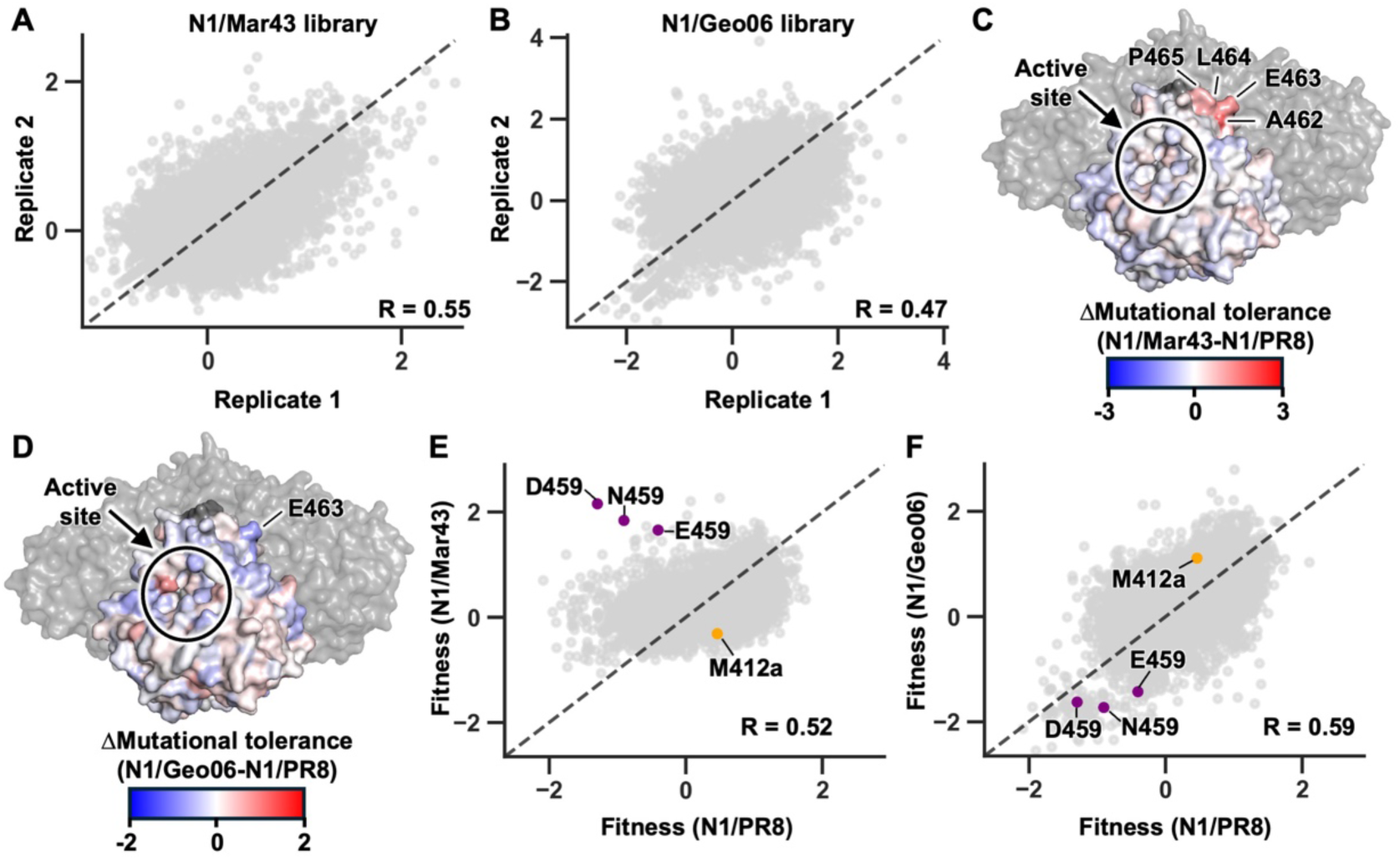
Reproducibility of the deep mutational scanning experiments and comparison of mutational effects between different H1N1 NAs. **(A and B)** Scatterplots showing the correlation of mutational fitness profiles between two biological replicates for (A) N1/Mar43 and **(B)** N1/Geo06. Each data point represents one mutation. The Pearson correlation coefficient (R) is indicated. **(C and D)** Difference of mutational tolerance between for **(C)** N1/Mar43 and N1/PR8 and **(D)** N1/Geo06 and N1/PR8 mapped onto the crystal structure of N1/Mar43 NA. One protomer is displayed in surface representation with residues colored according to mutational tolerance, while the remaining three protomers are shown in gray. Active site is circled. **(E and F)** Scatterplots showing the correlation of fitness scores for individual mutations between **(E)** N1/Mar43 and N1/PR8 and **(F)** N1/Geo06 and N1/PR8. H412aM, and P459D/N/E are indicated in orange and purple, respectively. Pearson correlation coefficient (R) is indicated.

**Figure S6.**
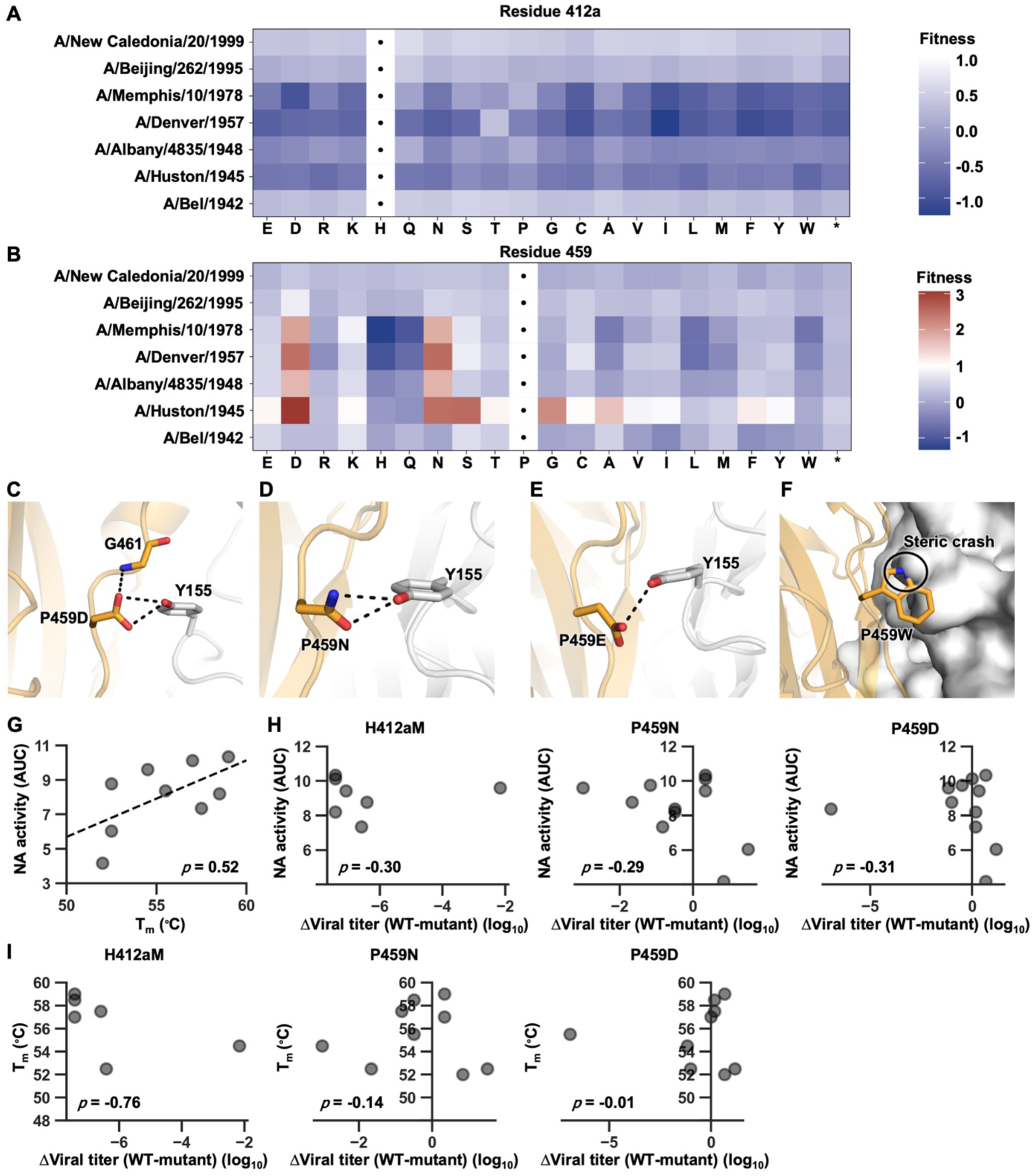
Mutational fitness effect at residues 412a and 459. **(A and B)** The fitness scores of mutations at **(A)** H412a and **(B)** P459 across different human H1N1 strains are shown as a heatmap. WT amino acids are indicated by a black circle. Stop codon is annotated as *. **(C-F)** Interactions between neighboring residues with P459D/N/E/W is shown. **(G)** Correlation between NA enzymatic activity and thermostability. NA activity was quantitated by area under curve (AUC). The temperature corresponding to the lowest point of the first derivative -d(RFU)/dT was determined to be the melting temperature (T_m_). Of note, thermostability data for A/Bel/1942 NA and N1/Mar43 NA are excluded from the analysis due to the presence of two distinct transitions in the thermal shift assay. **(H and I)** Correlation between **(H)** WT NA enzymatic activity or **(I)** thermostability and difference in viral titer between WT and H412aM or P459N/D. **(G-I)** The Spearman’s rank correlation coefficient (*ρ*) is indicated.

**Figure S7.**
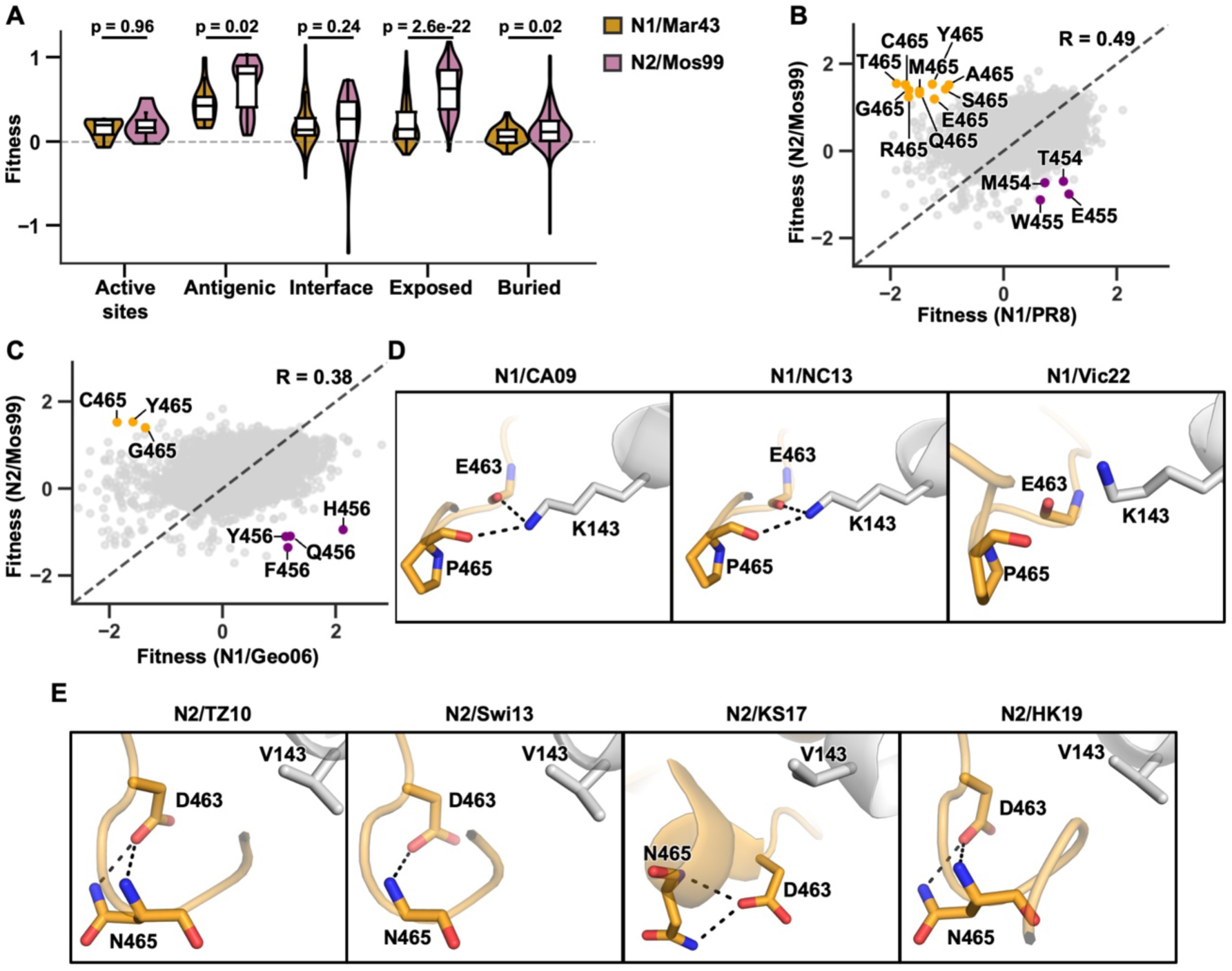
Comparison between N1 and N2 NAs. **(A)** Violin plots showing the distribution of mean fitness scores for active site, N1/N2 antigenic site, interface, exposed and buried residues. Yellow indicates the N1/Mar43 library, and pink indicates the N2/Mos99 library. Statistical significance was analyzed by two-sided Wilcoxon rank-sum tests, with p-values were adjusted using the Benjamini-Hochberg procedure. **(B and C)** Scatterplot showing the correlation of mutational fitness profiles between **(B)** N2/Mos99 and N1/PR8 as well as **(C)** N2/Mos99 and N1/Geo06. Each data point represents one mutation. Beneficial mutants in N2/Mos99 and N1/ PR8 or N1/Geo06 is indicated in orange and purple, respectively. Pearson correlation coefficient (R) is indicated. **(D)** Interactions between P465 and neighboring residues in different N1 are shown. H1N1 A/California/04/2009 (N1/CA09, PDB: 9O9V)^2^, H1N1 A/North Carolina/07/2013 (N1/NC13, PDB: 9EWQ)^3^, H1N1 A/Victoria/4897/2022 (N1/Vic22, PDB: 9DPC)^4^. **(E)** Interactions between N465 and neighboring residues in different N2 are shown. H3N2 A/Tanzania/205/2010 (N2/TZ10, PDB: 8G3Q)^5^, H3N2 A/Switzerland/9715293/2013 (N2/Swi13, PDB: 8ZYX)^6^, H3N2 A/Kansas/14/2017 (N2/KS17, PDB: 8ZR4)^7^ and H3N2 A/Hong Kong/2617/2019 (N2/HK19, PDB: 8G3V)^5^. Black dashed lines represent H-bonds.

